# CanSig: a tool for benchmarking malignant state discovery in single-cell RNA-Seq data

**DOI:** 10.1101/2022.04.14.488324

**Authors:** Florian Barkmann, Josephine Yates, Pawel Czyz, Agnieszka Kraft, Marc Glettig, Niko Beerenwinkel, Valentina Boeva

## Abstract

Single-cell RNA sequencing (scRNA-seq) facilitates the discovery of gene signatures that define cell states across patients, which could be used in patient stratification and drug discovery. However, the lack of standardization in computational methodologies to analyse these data impedes the reproducibility of signature detection. To address this, we developed CanSig, a comprehensive benchmarking tool that evaluates methods for identifying transcriptional signatures in cancer. CanSig integrates metrics for batch correction and biological signal conservation with a gene signature correlation metric to score according to rediscovery, cross-dataset reproducibility, and clinical relevance. We applied CanSig to ten methods and to ten scRNA-seq datasets from four human cancer types—glioblastoma, breast cancer, lung adenocarcinoma, and cutaneous squamous cell carcinoma— representing 116 patients and 105,000 malignant cells. Our results identify BBKNN as a leading method. We showed that the signatures identified with these methods correlate with clinically relevant outcomes, including patient survival and lymph node metastasis. Thus, CanSig establishes a standardized framework for reproducible cancer transcriptomics analysis.

## Introduction

For nearly a decade, single-cell RNA sequencing (scRNA-seq) has proven to be an essential tool for exploring the significant inter- and intra-patient heterogeneity found in malignant tissues [1,2]. While much of the research has centered on the heterogeneity within the tumor microenvironment, there has recently been growing evidence on the diversity of malignant cells both within and between patients. This has led to the identification of various gene signatures that represent unique cellular programs or states [3–11]. Understanding these cell states, including rare types like cancer stem cells, is crucial for cancer research, as they can significantly impact tumor maintenance, progression, and treatment resistance [2,12].

Despite progress, recent discoveries of gene signatures for shared transcriptional states in cancer have often lacked consistency across studies, resulting in signatures that are not reproducible or comparable across datasets. The methods used for signature discovery can generally be divided into two categories: early integration and late integration. Early integration methods merge data from all patients before identifying shared signatures in the combined dataset [13,14]. In contrast, late integration approaches identify signatures of differential states for individual patients and then aggregate them into shared signatures [3,4,10]. While early integration can enhance statistical power to detect rare transcriptional states that may be underrepresented in individual tumors, it risks introducing artifacts through the overcorrection of patient-specific effects. Late integration methods avoid this overcorrection but may be less effective in detecting rare cell states, which can sometimes be critical in cancer development [15].

Although existing early integration methods have been assessed in non-cancerous contexts, their applicability in cancer research remains uncertain. The integration of diverse populations of malignant cells presents unique challenges, such as accommodating variations in tumor genetic backgrounds [2,5], as well as addressing confounding factors like batch effects [16], sequencing depth [16], and cell cycle phases [5], which can obscure the biological signals of interest. Furthermore, there has been no comparative analysis of early versus late integration methods specifically for signature discovery, nor has the reproducibility of findings across different cancer types been explored.

In response to these gaps, we present CanSig, a benchmark designed to evaluate the discovery of shared transcriptional states in cancer cells. We assess various integration methods across multiple preprocessing conditions and their ability to rediscover established ground truth signatures in cancer cells, comparing both early and late integration strategies. Additionally, we evaluate the reproducibility and sample efficiency of the identified signatures, as well as their potential clinical relevance. Users can employ the CanSig framework complemented with tutorials and APIs on novel scRNA-seq datasets for *de novo* cancer state identification.

## Results

### CanSig benchmarks early and late integration strategies for signature discovery

CanSig provides a comprehensive benchmark of early and late integration strategies for the discovery of transcriptional signatures in single cancer cells. These signatures, which represent jointly activated pathways, provide insights into the biological functions of cells and are typically composed of sets of 50–200 co-expressed genes.

Signature discovery methods fall into two main categories: early and late integration (Fig. 1a). Early integration approaches identify a shared latent space across patients, correcting for batch effects, and then use clustering followed by differential gene expression analysis to derive shared transcriptional state signatures. A critical component of these methods is effective batch correction. In the CanSig benchmark, we evaluated integration methods such as ldvae [18], CCA [19], Combat [20], BBKNN [21], Harmony [22], Scanorama [23], Dhaka [24], and scVI [16] in terms of the batch correction efficiency and conservation of biological signal in the shared latent spaces, comparing the method performances to a baseline of no integration. Conversely, late integration methods first identify transcriptional programs within individual samples and subsequently aggregate these patient-specific programs into meta-signatures (Fig. 1a). We evaluated GeneNMF [25], a Python implementation of the commonly used non-negative matrix factorization (NMF), and scalop [7] as late integration strategies. Notably, only Dhaka and scalop were specifically designed to integrate cancer data, while the rest of the methods tested in CanSig were developed for the general purpose of single-cell data integration or signature discovery in non-mutated cells.

**Figure 1:**
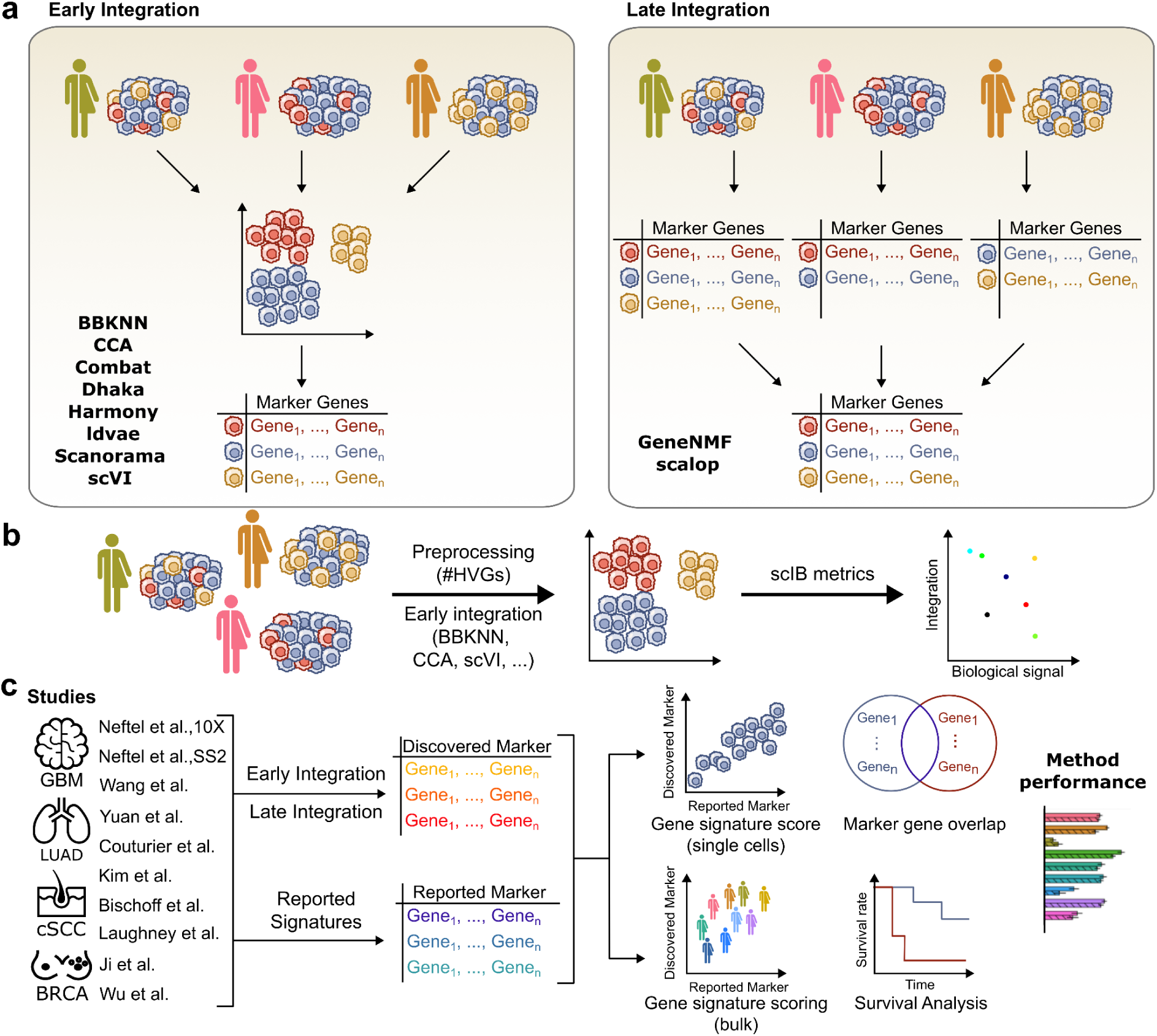
Overview of the CanSig benchmark. **a,** Comparison of early and late integration approaches for identifying novel shared transcriptional signatures in cancer. In early integration strategies, a common latent space is first established for cells from all samples in the study. This shared latent space is then used to identify signatures for shared transcriptional states, represented as lists of marker genes. In contrast, late integration strategies first identify transcriptional state signatures for each individual sample and then combine these into meta-signatures across patients. **b,** CanSig assesses integration methods in single-cell cancer datasets. After preprocessing samples, the effect of the strategy for selecting highly variable genes (HVG) on performance is evaluated. Data integration is performed using various methods, which are then assessed using scIB metrics [17], split into a batch correction (batch-corr) and a biological conservation score (bio-cons). The overall performance is calculated using the formula: *total score (TS) = 0.6 * bio-cons + 0.4 * batch-corr.* Thus, the top-performing methods must excel in both tasks, though preserving biological signals is given more weight than integration performance. **c,** Both early and late integration strategies are evaluated for their ability to rediscover previously identified transcriptional states in cancer studies, known as “ground truth signatures.” Early integration methods are used with Leiden clustering to identify signatures, and early and late integration performance is evaluated across three signature sets: the same number (*k*) as the ground truth signatures, *k*+1, and *k*+2 signatures. Rediscovery is assessed using a gene signature score (Methods) and marker gene overlap. Additionally, methods are evaluated for their capacity to discover clinically relevant signatures based on their performance in survival analysis and associations with clinical characteristics.

The first phase of the CanSig benchmark focused on evaluating batch integration methodologies independently of the signature discovery process (Fig. 1b). Datasets were processed using standard single-cell analysis workflows (Methods). A key preprocessing step involved selecting highly variable genes (HVG), and we assessed how variations in HVG selection—such as the number of HVGs and whether selection occurred at the patient level or across the full dataset—impacted method performance.

To facilitate cell labeling in the cancer setting, where annotation can be challenging, we used the maximum score of previously described signatures from the source studies as labels. These signatures, which have been validated in the literature, served as “ground truth” labels throughout the analysis. Integration performance was evaluated using two primary scores: a batch correction score (batch-corr) and a biological conservation score (bio-cons). The overall score was calculated as *TS = 0.6 * bio-cons + 0.4 * batch-corr*, placing greater emphasis on the preservation of biological signals over integration performance.

The second phase of the CanSig benchmark assessed both early and late integration strategies for their ability to rediscover “ground truth” transcriptional signatures (Fig. 1c). After applying Leiden clustering and differential gene expression analysis for early integration methods, we evaluated the top-performing early integration methods (BBKNN [21], CCA [19], scVI [16], and Harmony [22]), as well as late integration methods (GeneNMF [25] and scalop [7]), along with the baseline of no integration.

A commonly used measure for comparing signatures is marker gene overlap. However, this metric can be affected by differences in gene sets across datasets, even if the biological function is conserved. To address this, in addition to marker gene overlap, we introduced a gene signature scoring metric. This metric assesses the correlation between uncovered and ground truth signatures, penalizing cases where uncovered signatures correlate with too many ground truth signatures (Methods). This scoring was applied to both single-cell and bulk datasets, providing a more robust evaluation of signature similarity.

Lastly, transcriptional programs identified in single cancer cells are often used to infer biological functions and clinical characteristics. Programs with strong clinical associations are particularly valuable for their potential clinical applications. Therefore, we evaluated methods based on their associations with survival, lymph node metastasis, and therapy response, assessing their ability to stratify patients effectively.

### Early integration strategies show variable performance on single cancer cell datasets

We assessed eight integration strategies across two glioblastoma (Neftel *et al.,* 10X and Smartseq2) [7], one lung adenocarcinoma (Kim *et al.*) [26], and one cutaneous squamous cell carcinoma (Ji *et al.*) [9] dataset (Fig. 2a). Certain integration methods, such as Combat and Dhaka, consistently underperformed in both batch correction and biological conservation, while others, like ldvae, struggled primarily with batch integration. Interestingly, the baseline of no integration often outperformed several methods in preserving biological signals.

**Figure 2:**
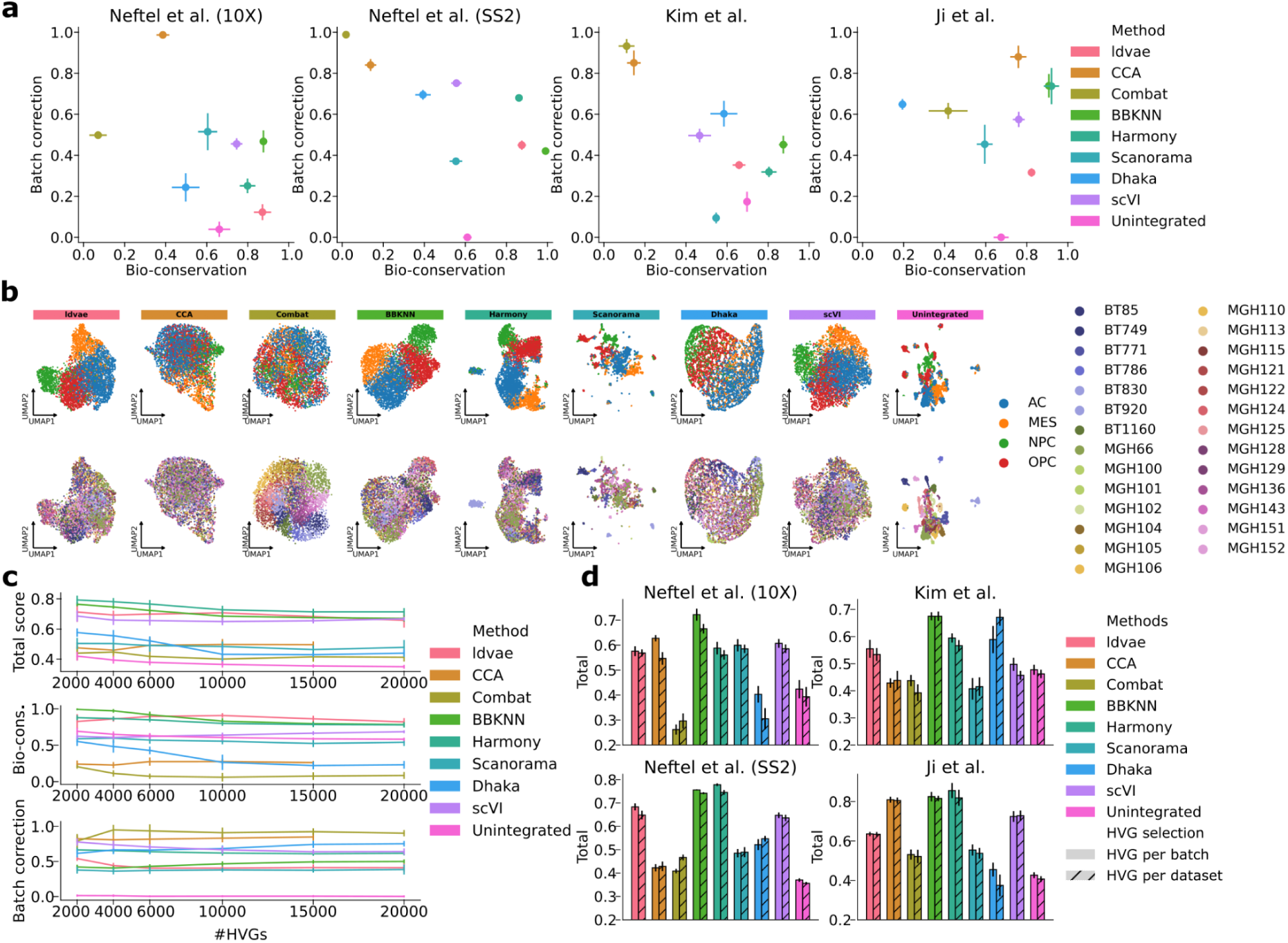
CanSig evaluates early integration strategies. **a**, scIB metrics [17] scores calculated for 8 early integration strategies [16,18–24], with unintegrated data serving as a baseline. The methods were applied to the Neftel et al. glioblastoma dataset [7], split by sequencing technology (10X and SmartSeq2); the Kim et al. lung adenocarcinoma dataset [26]; and the Ji et al. cutaneous squamous cell carcinoma dataset [9]. On the x-axis, the bio-conservation score is shown, which integrates metrics based on both cell label-dependent and label-free conservation. The y-axis represents the batch correction score, which combines metrics with and without reliance on cell label identity (Methods) [17]. Cell labels are assigned using the maximum score of signatures from the original publications (Methods). Each method was run on five subsampled sets, with 80% of patients randomly selected, to estimate the median score (represented by dots) and 95% confidence interval for each method (represented by bars). **b,** UMAP plots for the latent spaces generated by each method for the Neftel SmartSeq2 dataset [7]. Cell labels are assigned as in panel **a**. The top row shows cell labels, and the bottom row shows patient IDs. **c,** The evolution of the total score, biological conservation score, and batch correction score as a function of the number of highly variable genes selected for each method. Notably, CCA could not run with more than 15,000 genes. **d,** The impact of selecting highly variable genes at the batch level (HVG per batch) versus across the entire dataset (HVG per dataset) on the total score. The 95% confidence interval (CI) across subsampled runs is represented by bars.

In contrast, methods like BBKNN and Harmony demonstrated robust performance across datasets. This is further illustrated in the low-dimensional cell embeddings (Fig. 2b), where BBKNN and Harmony spaces show clear separation of cell types and strong mixing of cells coming different patients. In comparison, Combat and Dhaka exhibited poor cell type separation and visible clustering of patient-specific cells. Overall, BBKNN consistently outperformed or performed on par with other integration methods in both biological conservation and batch correction in cancer datasets.

### Strategies for the selection of highly variable genes before integration have little influence on integration performance

We next assessed how the selection of highly variable genes (HVGs) for single-cell intergration influenced biological preservation, integration, and overall performance scores. First, we examined how increasing the number of HVGs affected these scores (Fig. 2c). Conceptually, an optimal range of HVGs should exist, as a smaller set may capture insufficient information, while a larger set could introduce excessive noise and increase computational demands. Across all methods, except for CCA, we observed that the total score either decreased slightly or remained stable as more HVGs were added. This decline was primarily reflected in the biological conservation score, with the integration score remaining relatively constant. Our analysis suggests that selecting 2,000–4,000 HVGs is generally preferable, although this parameter had only a minor effect on overall performance.

We also evaluated the impact of patient-specific versus full dataset HVG selection on method performance (Fig. 2d). In this approach, HVGs were selected separately for each patient and then aggregated, preventing larger patient samples from disproportionately influencing the analysis. We found that patient-specific HVG selection generally improved performance for most methods. Notably, for the top-performing methods, BBKNN and Harmony, patient-specific HVG selection consistently outperformed full-dataset selection, though not always significantly.

In summary, the HVG selection process had a modest effect on integration performance, with optimal results achieved by selecting a lower number of HVGs and employing patient-specific HVG selection.

### Early and late integration methodologies perform variably on signature rediscovery

To assess the ability of early and late integration strategies to uncover *de novo* shared transcriptional states in single cancer cells, we evaluated their performance in rediscovering ground truth signatures across various cancer types and datasets, including an additional breast cancer dataset from Wu et al. [10].

BBKNN once again emerged as a top-performing method, consistently rediscovering signatures across different cancer types and datasets (Fig 3a). It showed better average score across datasets across all settings (using the true number of signatures (*k*) as input, *k+1*, and *k+2*), It outperformed other methods on the Ji et al. (3), Neftel et al. 10X (4), and Neftel et al. 10X (6) datasets, and ranked second on the Neftel et al. SS2 (4) and Kim et al. (3) datasets. Some methods showed considerable variability across datasets; for example, scalop performed exceptionally well on the Neftel et al. SS2 (4) and SS2 (6) datasets, where it was originally developed, but underperformed in other contexts, such as Ji et al. (3), Neftel et al. 10X (6), and Kim et al. (3). Notably, no method successfully rediscovered the original signatures reported in the Wu et al. breast cancer dataset, with the best methods achieving only an average score of ∼0.4.

**Figure 3:**
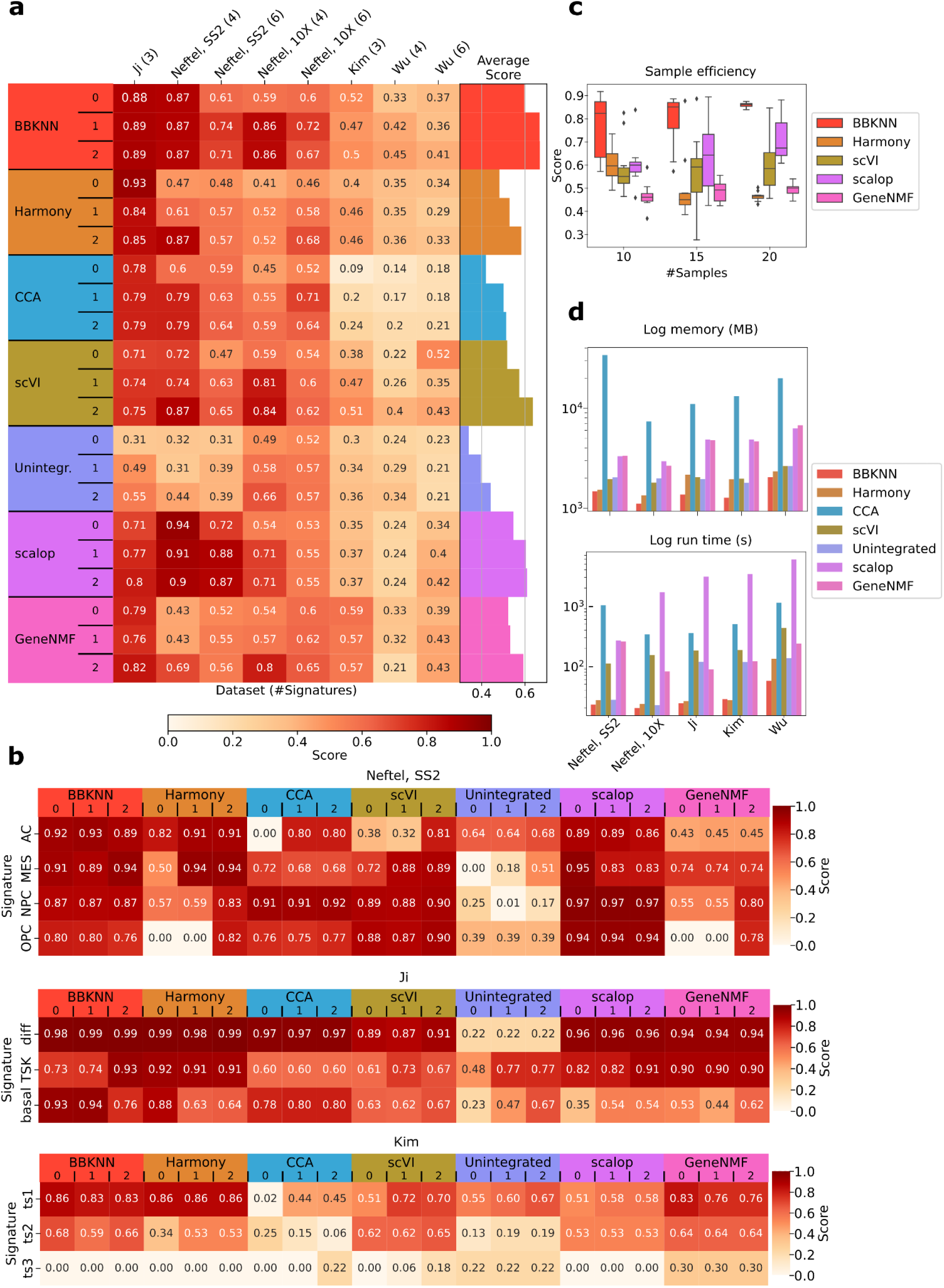
Evaluation of early and late integration strategies for signature rediscovery. **a,** CanSig benchmarks early and late integration methods across multiple cancer datasets, including the previously introduced datasets and an additional breast cancer dataset from Wu et al. [27], to assess cross-dataset signature rediscovery. The y-axis indicates the datasets, with the number of originally reported signatures shown next to each dataset’s name. In the glioblastoma dataset, the four primary states were subdivided into six states, and methods were evaluated for rediscovery in both scenarios. For early integration strategies, the methods were run using the true number of signatures (*k*), *k*+1, and *k*+2 as input parameters for Leiden clustering, while late integration strategies were evaluated with these same signature counts for the entire method. These conditions are denoted in the rows as 0, 1, and 2, respectively. The heatmap shows the single-cell gene signature score (Methods), with higher scores reflecting better rediscovery of ground truth signatures. The barplots on the right represent the average score across datasets of a specific run. **b,** Cancer type-specific rediscovery scores for glioblastoma (Neftel et al. SS2), lung adenocarcinoma (Kim et al.), and cutaneous squamous cell carcinoma (Ji et al.). Heatmaps follow the same layout as in panel **a,** except that scores are displayed individually for each ground truth signature rather than averaged across them. **c,** Efficiency of early and late integration methods as a function of sample size demonstrated by subsampling the Neftel et al. SS2 dataset to 10, 15, and 20 patients out of 27 with the total gene signature score for the four-state ground truth signatures with four clusters reported. **d,** Memory and time requirements of the evaluated methods across datasets. The log memory (in MB) and log run time (in s) is indicated for each dataset and each method.

### Early integration methods are more sample-efficient

Since early integration methods initially integrate cells across all patients, they may more readily rediscover signatures in datasets with fewer samples. To test this hypothesis, we downsampled the Neftel et al. SS2 dataset to 10, 15, and 20 samples and evaluated the gene signature scores for three early integration methods (BBKNN, Harmony, and scVI) and two late integration methods (GeneNMF and scalop). Results showed that early integration methods like BBKNN maintained strong performance across all sample sizes, whereas late integration methods significantly improved with more samples available (Fig. 3c). These findings suggest that early integration methods, particularly BBKNN, are better suited for settings with limited sample numbers, maintaining stable performance across sample sizes.

### Early integration methods are more memory and time efficient

We assessed the computational performance of various methods across datasets, focusing on memory and runtime requirements. Among all methods evaluated, BBKNN consistently demonstrated the lowest memory usage and fastest execution times, making it the most computationally efficient option.

In general, early integration methods required significantly less memory compared to late integration strategies, with the exception of CCA, which exhibited higher memory demands. Similarly, early integration methods outperformed in terms of runtime, with BBKNN and Harmony emerging as the fastest methods across all datasets. In contrast, scalop required the longest runtime in four out of five datasets analyzed.

Overall, early integration methods, particularly BBKNN and Harmony, exhibited markedly lower computational demands than late integration approaches, enhancing their practical usability.

### Most methods struggle to rediscover gene signatures across datasets

To assess whether signatures identified in one dataset are reproducible in external datasets—a task rarely addressed by studies reporting “ground truth” signatures—we evaluated the gene signature scores in three independent GBM datasets (Yuan *et al*. [6], Couturier *et al*. [30], and Wang *et al*. [28]) and two external LUAD datasets (Bischoff *et al*. [29], Laughney *et al*. [31]).

Our analysis revealed that early integration methods such as BBKNN and scVI were the most effective in rediscovering the original signatures across external datasets, followed by late integration strategy scalop (Fig. 4a). In particular, these methods rediscovered the four Neftel *et al*. signatures across all three GBM datasets the best. Notably, performance differences often stemmed from specific signatures being poorly recovered; for instance, CCA, Harmony, and BBKNN struggled with the OPC signatures in the Yuan *et al*. dataset and the MES signature in the Wang *et al.* dataset, while scalop and scVI struggled with the NPC signature in the Wang *et al.* dataset. (Fig. 4b).

**Figure 4:**
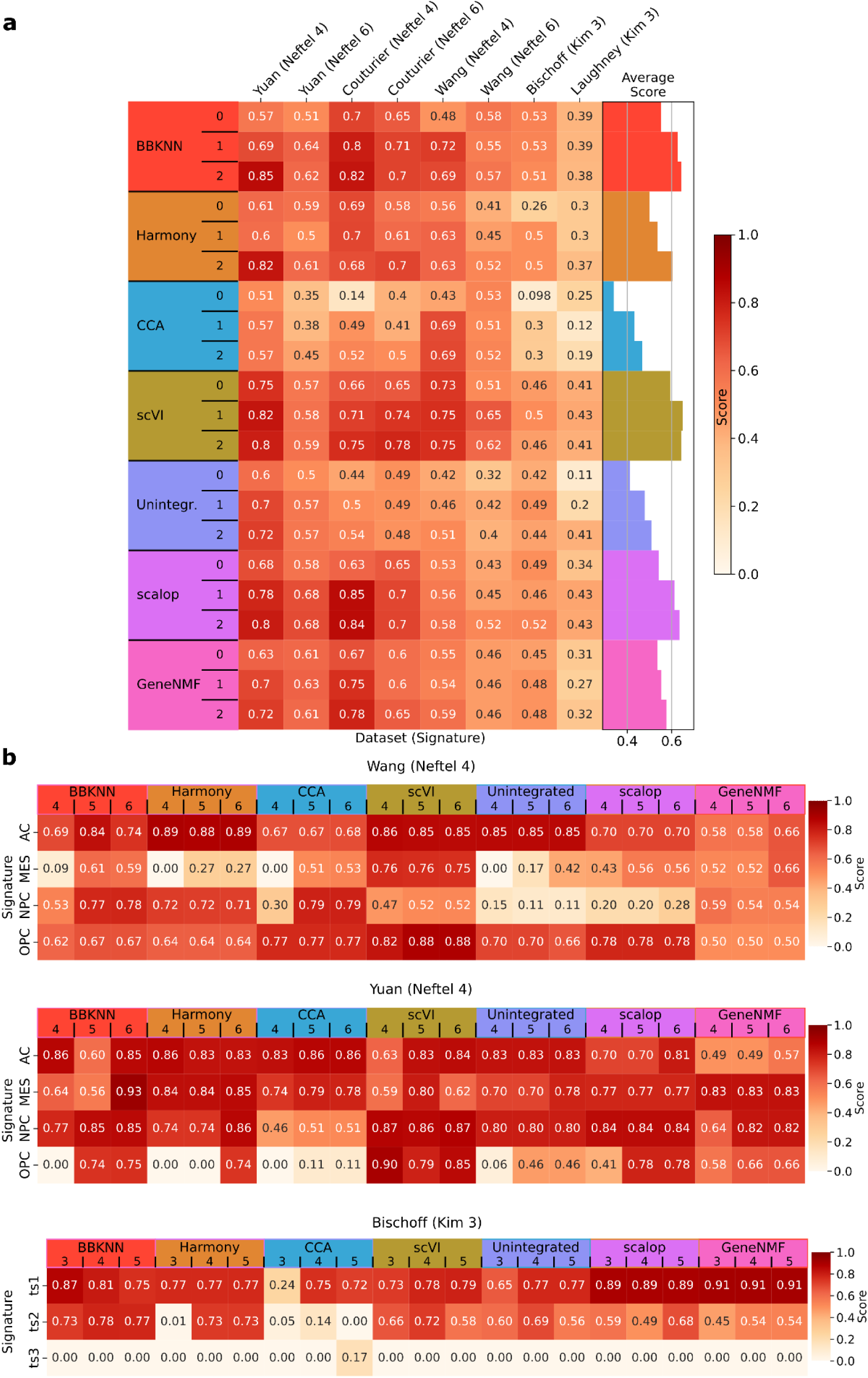
Evaluation of reproducibility of signature discovery across datasets. **a**, Heatmap comparison of signature rediscovery performance across multiple cancer datasets. Columns show different datasets (with original signature counts noted). **b,** Scores for individual signatures on new datasets: a glioblastoma dataset from Wang et al. [28], Yuan *et al*. [6], and a lung adenocarcinoma from Bischoff *et al*. [29]. Heatmaps follow the same layout as in panel **a,** except that scores are displayed individually for each ground truth signature rather than averaged across them.

All methods exhibited reduced performance in the GBM setting when evaluating six signatures, with no method achieving high rediscovery scores in the Wang *et al*. dataset. Similarly, the LUAD signatures were challenging to reproduce across methods, with the highest scores (>0.5) observed for BBKNN and scalop (Fig. 5a). The poor rediscovery was particularly pronounced for the signature of the ts3 state, which none of the methods could accurately capture in the Bischoff *et al.* dataset. This signature, as originally described, corresponds to a rare transcriptional state resembling normal ciliated cells, underscoring the difficulty of replicating certain signatures across datasets (Fig. 5b).

**Figure 5:**
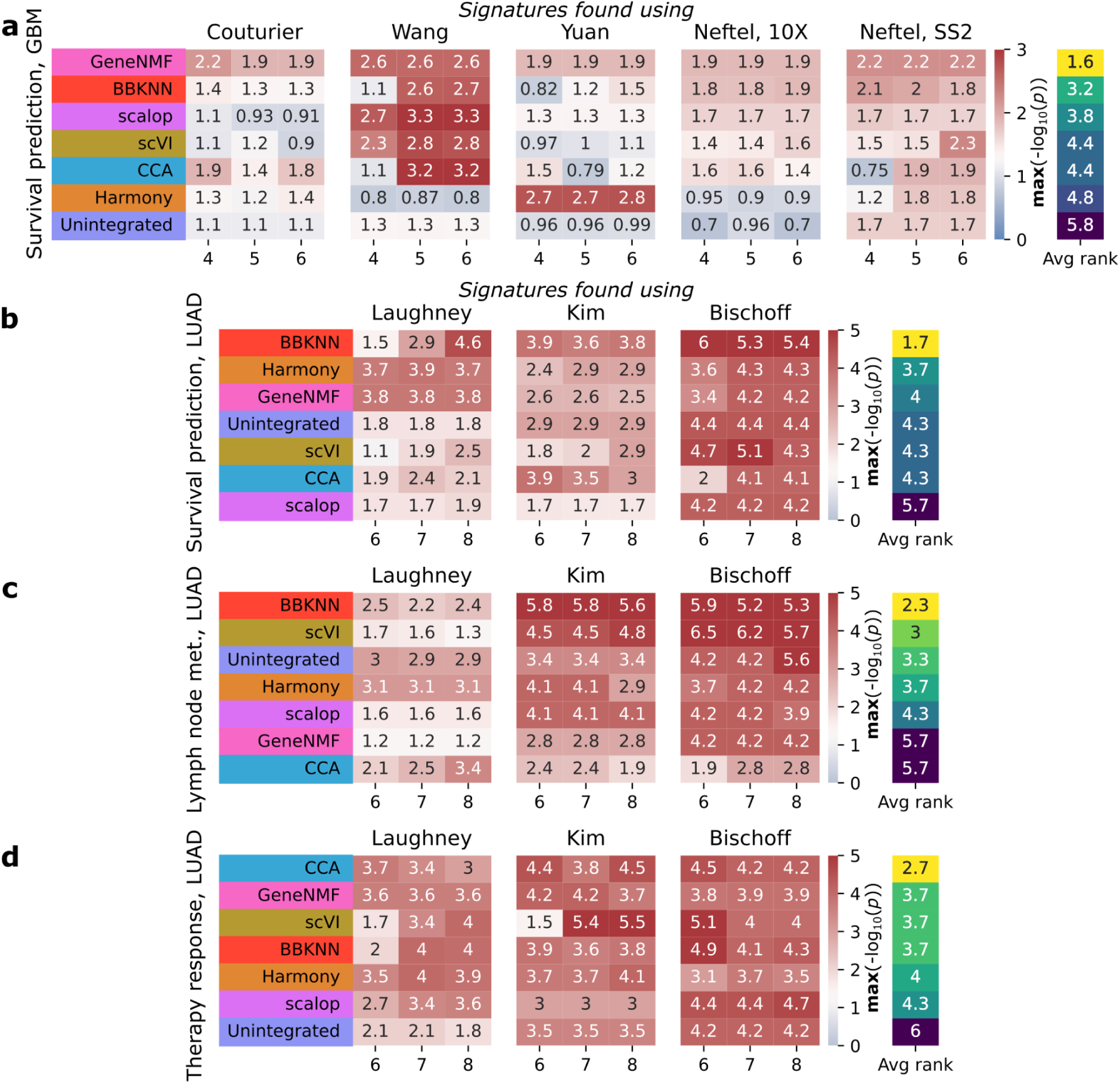
Association between discovered signatures and survival and clinical features across methods. **a-b,** Association between signatures uncovered through a range of methods and survival in **a**, the TCGA glioblastoma (GBM) and **b**, lung adenocarcinoma (LUAD) dataset. Signatures are uncovered in single-cell data of glioblastoma [6,7,28,30] and lung adenocarcinoma datasets [26,29,31], respectively. Then, these signatures are scored in bulk TCGA data of the corresponding cancer type. For each uncovered signature, a univariate Cox model is fitted to predict survival using signature scores and obtain the p-value associated with the beta coefficient of the signature. Rows indicate the method used to discover signatures and the columns represent the number of signatures discovered. Each entry indicates the maximal -log_10_(p-value) across all signatures. Methods are ranked for each dataset according to the average maximal -log_10_(p-value) across different numbers of signatures; the average rank across datasets is then used to obtain a final ranking of all signatures, indicated in the last column. **c-d,** Association between signatures uncovered through a range of methods in lung adenocarcinoma datasets and **c**, lymph node metastasis and **d**, therapy response. Patients are classified into two categories, either having any lymph node metastasis present or none for **c**, or showing progressive disease as opposed to stable, partial, or complete remission for **d**. The TCGA signature scores are then compared between the two groups using a Kruskal-Wallis test. As in **a-b**, the rows represent methods and columns the number of signatures, whereas each entry in the heatmap indicates the maximal -log_10_(p-value) across all signatures of association with lymph node metastasis (resp. therapy response).

These findings highlight the challenges of signature reproducibility across varying technologies and batch effects, emphasizing the need for robust cross-dataset validation. Overall, BBKNN, scVI, and scalop consistently demonstrated competitive performance in this setting.

### BBKNN outperforms other methods in discovering signatures associated with survival and clinical characteristics

Identified programs in cancer cells serve important roles in downstream analyses aimed at elucidating biological and clinical characteristics. A key application of these malignant program signatures is patient stratification based on survival outcomes. Patients are evaluated based on the proportion of tumor cells expressing these identified programs and these scores are used to make survival predictions. Additionally, these signatures can be linked to clinical features, such as lymph node metastasis and therapeutic response. Signatures that demonstrate stronger predictive capabilities for survival or closer associations with clinical characteristics potentially possess greater clinical utility as they can enable improved patient stratification and the formulation of new hypotheses related to tumorigenesis, metastatic behavior, and therapy resistance. Consequently, methodologies that yield signatures with stronger correlations to survival and clinical outcomes are particularly valuable.

To evaluate the relationship between the identified signatures and survival as well as clinical characteristics, we conducted an assessment using external bulk datasets obtained from The Cancer Genome Atlas (TCGA). For survival analysis, univariate Cox models were employed to investigate the association between scores of each signature and overall survival (Methods, Fig. 5a-b). Our findings indicated that in glioblastoma (GBM) and lung adenocarcinoma (LUAD), the efficacy of various methods in stratifying patients considerably differed. For instance, within the GBM cohort, none of the signatures derived from the Couturier *et al.* dataset [30] using scVI exhibited significant predictive power (*p* = 0.06), while the most predictive signature generated via GeneNMF demonstrated a strong association with survival (*p* = 0.006) (Fig. 5a). In LUAD, the most predictive signature generated from the Kim *et al.* dataset [26] revealed a slight association with survival when derived from scalop (*p* = 0.02), whereas the BBKNN signature showed a highly significant predictive capacity (*p* = 0.0001) (Fig. 5b). To systematically evaluate the predictive capacity of the methodologies, we ranked them based on the average minimum *p*-value across datasets (Methods). Notably, BBKNN consistently emerged as the most reliable method for uncovering predictive signatures, with an average rank of 3.2 in glioblastoma (second to GeneNMF) and an average rank of 1.7 in lung adenocarcinoma.

Furthermore, we assessed the association of the identified signatures with clinical characteristics, particularly the presence of lymph node metastasis and therapy response in LUAD (Methods; Fig. 5c-d). It is important to note that data regarding these clinical features were unavailable for the GBM cohort. Again, the methods displayed varying capabilities in identifying signatures linked to these characteristics; for example, the most associated of CCA’s signatures found using the Kim et al. dataset associated more weakly with lymph node metastasis (p=0.004) than BBKNN that found a highly significantly associated signature (p=1.6e-6) (Fig. 5c). Similar to the survival analysis, we ranked the methodologies based on the average minimum *p*-value across datasets. BBKNN emerged as one of the best performing methodologies, with an average rank of 2.7 for lymph node metastasis association (making it the best ranking method before scVI) and an average rank of 3.7 on therapy response association, 3-way tied in second place with GeneNMF and scVI (Fig. 5d).

In summary, signatures derived from different methodologies exhibited varying degrees of clinical and biological relevance. In line with the overall good and stable performance of BBKNN in previous comparisons, these results demonstrated the clinical relevance of the BBKNN-derived signatures of transcriptional states in cancer; signatures obtained through this approach consistently correlated with survival outcomes in GBM and LUAD, as well as with lymph node metastasis and therapy response in LUAD.

## Discussion

In this work, we systematically evaluated early and late integration strategies to identify optimal approaches for uncovering novel, shared transcriptional states in single cancer cells. Although recent studies have reported shared transcriptional states across cancer types, methodological differences make some findings challenging to reproduce across datasets, potentially compromising biological reliability. Here, we benchmarked eight early integration methods to assess their batch-correction capability and ability to retain biological signal, finding BBKNN [21] and Harmony [22] as consistent top performers. While highly variable gene (HVG) selection showed minimal effect on performance, selecting 4,000 HVGs in a batch-specific manner yielded the best outcomes.

In comparing early and late integration strategies on signature rediscovery tasks, early integration methods, particularly BBKNN, demonstrated strong performance across cancer types and conditions. While no method excelled in cross-dataset signature rediscovery, early integration approaches, including BBKNN, proved more sample-efficient, performing well on smaller datasets. Additionally, some methods better identified signatures associated with clinical features such as patient stratification based on survival, treatment response, and lymph node metastasis status.

Overall, BBKNN appears to be a robust choice for identifying shared transcriptional states across datasets, while early integration methods generally offer advantages on smaller datasets (n<10), in contrast to late integration strategies, which may be more effective on larger datasets.

Our analysis has some limitations. First, while we utilized well-validated, literature-reproduced signatures as “ground truth” states, assigning malignant cells to signatures remains challenging, making cell-type annotations potentially less accurate in this setting. Second, cross-dataset rediscovery was limited to two cancer types, as few datasets with reliable transcriptional states were available, which may limit the study’s generalizability. Finally, while we assess methods based on their ability to uncover clinically meaningful states, it is worth noting that states highly correlated with clinical factors, such as disease stage, may not necessarily reflect novel biological discoveries.

This work represents the first comprehensive assessment of integration strategies specifically in a cancer context, highlighting the unique challenges and providing recommendations for robust cancer transcriptional state discovery. We anticipate this study will guide the development of reproducible, clinically relevant cancer cell state signatures and foster more consistent findings across cancer research.

## Methods

### Datasets

Ten studies were used in this work: five glioblastoma datasets, comprising studies by Neftel *et al*. [7] (split by sequencing technology, 10X and SmartSeq2), Wang *et al*. [28], Yuan *et al*. [6] (subsetted to IDH-mutant GBM) and Couturier *et al*. [30]; three lung adenocarcinoma datasets, comprising studies by Kim *et al*. [26], Bischoff *et al*. [29], and Laughney *et al*. [31]; one squamous cell carcinoma dataset from Ji *et al*. [9] and one breast carcinoma dataset from Wu et al.[10]. These cancer types were chosen because the original studies reported distinct cancer cell states and associated gene signatures. For glioblastoma datasets, we used four primary signatures from the Neftel et al. work [7]: astrocyte-like (AC), mesenchymal-like (MES), neural progenitor cell-like (NPC), and oligodendrocyte-like (OC). Additionally, as the MES and NPC states were subdivided by the original authors into MES1/MES2 and NPC1/NPC2, we also performed analyses using this six-signature set. For squamous cell carcinoma, we utilized three signatures reported in the Ji et al. work [9]: differentiated, basal, and tumor-specific keratinocyte (TSK). Lastly, in lung adenocarcinoma datasets, we applied three state-specific signatures derived from the Kim et al. paper [26]: ts1, ts2, and ts3.

### Data preprocessing

For data preprocessing, we followed the standard processing guidelines described at https://www.sc-best-practices.org/preprocessing_visualization/quality_control.html. For datasets available only in transcripts per million (TPM), such as the Neftel et al. dataset generated with SmartSeq-2 (SS2), we converted TPM values to approximate raw counts by rounding to the nearest integer, as certain methods (e.g., scVI) require raw count inputs. Next, we filtered the datasets to include only malignant cells, and removed cells with less than 1000-3000 genes expressed, 1500 transcripts, and more than 20-30-% mitochondrial counts, with thresholds adjusted per sequencing technology and dataset. Cells were scored for cell cycle phase, retaining only those in G1. Samples with fewer than 50 remaining cells were removed, and genes expressed in less than 0.1% of remaining cells were filtered out. Finally, we normalized the data using TPM values, followed by log1p transformation. We assigned cell states to malignant cells using the maximum scores after Scanpy scoring for the previously validated “ground truth” signatures reported in the literature.

### Early integration strategy evaluation

We evaluated 8 early integration strategies [16,18–24], with unintegrated data serving as a baseline. These methods were selected based on their prevalence within the community (BBKNN, Harmony, Scanorama, scVI, CCA), their use of simpler models for batch correction (Combat, ldvae), or their design for integrating cancer data (Dhaka).

– CCA [19] integrates single-cell datasets from multiple conditions or technologies using canonical correlation analysis. It identifies shared canonical correlation vectors across datasets and mutual nearest neighbors, capturing common sources of variation while allowing for dataset-specific differences.
– Combat [20] corrects batch effects in RNA-seq count data using a negative binomial regression framework, adjusting for batch-specific differences without altering biological variation.
– BBKNN [21] builds a batch-aware k-nearest neighbors graph to correct batch effects in single-cell data, ensuring that neighbors are balanced across batches while preserving the local data structure.
– ldvae [18] utilizes a variational autoencoder (VAE) to identify latent factors in single-cell RNA-seq data, incorporating interpretability constraints like sparsity and biological priors to enhance the biological relevance of the latent representations.
– Harmony [22] corrects for batch effects by iteratively adjusting a low-dimensional embedding, maximizing cell mixing across batches while preserving biological variation through an alternating optimization strategy.
– Scanorama [23] integrates datasets by identifying and aligning shared features through manifold matching, using an efficient algorithm to iteratively merge datasets into a unified representation.
– scVI [16] employs conditional variational autoencoders (CVAEs) to capture the latent structure of single-cell data, explicitly modeling gene expression distributions and accounting for noise and technical variation.
– Dhaka [24] applies a specialized VAE to model and dissect tumor heterogeneity in single-cell genomic data, using a zero-inflated negative binomial distribution to address sparsity and overdispersion, and making it the only early integration method designed specifically for cancer data.

The methods were applied to the Neftel et al. glioblastoma datasets [7], split by sequencing technology (10X and SmartSeq2); the Kim et al. lung adenocarcinoma dataset [26]; and the Ji et al. cutaneous squamous cell carcinoma dataset [9]. By default, each method was run using a selection of 4,000 highly variable genes (HVGs) determined by the per-batch selection strategy in Scanpy.

Integration performance was assessed using two main scoring categories from the scIB benchmark [17]: a batch correction score (batch-corr) and a biological conservation score (bio-cons). The biological conservation score combines several metrics that leverage ground truth labels—specifically, normalized mutual information (NMI), adjusted rand index (ARI), average cell-type silhouette width, graph cell-type local inverse Simpson’s Index (cLISI), isolated label F1, and isolated label silhouette. The batch correction score incorporates label-based metrics such as principal component regression, average batch silhouette width, graph connectivity, graph integration local inverse Simpson’s Index (iLISI), and the k-nearest neighbor batch effect test (kBET).

To assess variability, we subsampled the datasets to 80% of patients five times, running each method on each subsampled dataset to estimate mean performance and standard deviation across iterations.

To assess the impact of HVG selection on performance, we varied the number of highly variable genes selected and measured the resulting performance across all methods on the Neftel et al. SS2 dataset. Additionally, we examined performance differences when HVGs were selected per batch compared to selection across the entire dataset, conducting this analysis on all four datasets.

### Early and late integration strategy evaluation

The primary objective of these methods is to identify transcriptional signatures representing distinct programs or states in malignant cells. To evaluate different methods, we assessed each method’s ability to rediscover predefined “ground truth” signatures. For early integration methods, we generated signatures by applying Leiden clustering and differential gene expression analysis to the latent spaces produced, whereas late integration methods inherently yielded signatures. We assessed the highest-performing early integration methods (BBKNN [21], CCA [19], scVI [16], and Harmony [22]), along with the late integration methods (GeneNMF [25,32] and scalop [7]) and included an unintegrated baseline. We used these two integration methods are they represent the two main variations of late integration used in the field, either non-negative matrix factorization (NMF)-based or clustering-based.

– GeneNMF [25,33] applies NMF to multiple samples to discover gene programs and identify consensus patterns, termed meta-programs (MPs), across these samples. It clusters similar gene programs across datasets, calculates MP signature scores, and identifies differentially expressed genes.
– scalop [7] builds a dendrogram using average linkage and Pearson correlation to cluster malignant cells in each sample based on gene expression, then filters clusters based on criteria such as cell count, gene expression differences, and statistical significance. It then refines the clusters by removing redundant or cell cycle-related signatures, and clusters the remaining signatures into meta-modules, selecting genes with significant expression changes as signature genes for further analysis.

For evaluation, we first used marker gene overlap, a common measure for signature rediscovery. However, this metric has limitations, especially when dataset-specific gene sets vary while underlying biological functions remain consistent. To address this, we introduced an additional gene signature scoring metric that leverages signature scores to compare the expression profiles of discovered and ground truth signatures. Instead of directly matching gene sets, this metric assesses whether the two signatures are expressed in the same cells, indicating shared biological activity, thus producing high correlation scores across cells when biological functions are indeed similar.

Let ρ be the Spearman correlation, ν_*i*_ the score of the i-th ground truth signature, η_*i*_ the score of the i-th signature uncovered by the method, and 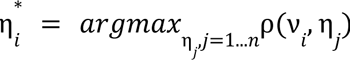.

The gene signature score is

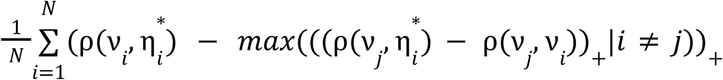

The first component of the scoring metric calculates the Spearman correlation between the ground truth signature and the most highly correlated uncovered signature. The second component applies a penalty to the uncovered signature if it exhibits strong correlation with additional ground truth signatures, reducing its score by the degree of overlap between the original ground truth signatures. Only the positive portion of this result is retained, as negative scores are not desired. This scoring approach was applied to both single-cell and bulk datasets. For single-cell data, individual cells were scored using Scanpy’s scoring function, while for bulk datasets, the average Z-score of marker genes was used, where the Z-score of gene i in patient j is 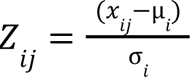, with *x_ij_* the original expression of the gene, µ the average of the gene over all patients, and σ the standard deviation of the gene over all patients.

To assess performance across varying numbers of uncovered signatures, each method was initially run with the number of signatures *k* matching the ground truth signature count. Recognizing that the exact number of transcriptional programs or cell states is often unknown, we additionally ran each method with one (*k+1*) and two (*k+2*) extra signatures. Methods were evaluated across this range for each dataset. For the glioblastoma datasets specifically, evaluations were conducted using both four and six ground truth signatures.

### Cross-dataset signature discovery evaluation

To evaluate the ability of different methods to rediscover previously reported signatures in new datasets, we tested their performance in identifying Neftel et al.’s [7] “ground truth” signatures across three external glioblastoma datasets (Yuan et al. [6], Wang et al. [28], and Couturier et al. [30]) and Kim et al.’s [26] “ground truth” signatures across two external lung adenocarcinoma datasets (Laughney et al. [31] and Bischoff et al. [29]). The evaluation followed the same approach outlined in the “Early and Late Integration Strategy Evaluation” section.

### Sample efficiency analysis

To investigate whether early integration strategies are better suited for capturing “ground truth” signatures in low-sample scenarios, we compared the performance of three early integration methods (BBKNN [21], scVI [16], and Harmony [22]) with two late integration methods (scalop [7] and GeneNMF [25,32]). This analysis was conducted on Neftel et al.’s Smart-seq2 dataset [7], downsampled to include 10, 15, and 20 samples out of the original 27. For each subset, we calculated the total gene signature score associated with the rediscovery of Neftel et al.’s four reported signatures.

### Link between uncovered signatures and survival and clinical characteristics

To evaluate how associated the uncovered signatures are with survival and clinical characteristics, we first scored the signatures uncovered in the single-cell data in external bulk datasets. We selected bulk datasets in the Cancer Genome Atlas that corresponded to the original cancer types in which the signatures were discovered: glioblastoma (GBM) for the Neftel et al. [7], Wang et al. [28], Couturier et al. [30], and Yuan et al. [6] datasets; lung adenocarcinoma (LUAD) for the Kim et al. [26], Laughney et al. [31], and Bischoff et al. [29] datasets. Unfortunately, no bulk dataset corresponded to the Ji et al. [9] dataset, so we did not evaluate signatures uncovered in this single-cell dataset.

Signatures are named according to the single-cell dataset from which they were derived (e.g., Kim et al.). To score TCGA patients, we calculated the average Z-score of marker genes for each signature, as previously described.

For survival analysis, these scores were used as input to a univariate Cox proportional hazards model, implemented using the lifelines package [33]. This model provided the beta-coefficient for each score, along with its p-value determined by a t-test.

For associations with clinical characteristics, we compared signature distributions across patient groups using a Mann-Whitney U test. Clinical categorizations included lymph node metastasis status (no metastasis = 0, any metastasis = 1) and disease progression (progressive disease = 1, complete/partial remission or stable disease = 0). For each task, the maximum -log10(p-value) across all signatures is reported.

## Code and data accessibility

The CanSig code is accessible at https://github.com/BoevaLab/CanSig-benchmark.

